# *MITScan*: Score-Based Genome-Wide Association Analysis of the Microbiome and Host Transcriptome

**DOI:** 10.1101/2025.06.24.661342

**Authors:** Yajing Hao, Tal Kafri, Fei Zou

## Abstract

Microbiome research explores how microbial communities interact with the human body to influence health and well-being, offering insights for advancing personalized medicine and targeted treatments. Microbiome profiles can be measured, for example, by 16S rRNA sequencing, where sequencing reads are clustered into genus-or species-level operational taxonomic units (OTUs), yielding count-based abundance data. In addition, these data are often characterized by sparsity with extra zeros.

Emerging evidence suggests that imbalances in microbial composition, potentially regulated by host gene expression, have been associated with various diseases. Investigating the associations between the microbiome and host transcriptome can help uncover the mechanisms underlying human health and disease. Current existing approaches for evaluating their associations largely rely on Pearson or Spearman correlations, ignoring the fact that microbiome data are sparse count data, leading to potentially biased results. To overcome this issue, we present *MITScan*, a genome-wide ***MI*** crobiome-host ***T*** ranscriptome ***Sc***ore-based association ***an***alysis, employing zero-inflated negative binomial models to accommodate extra zeros commonly observed in microbiome data. To address the large number of paired association tests arising from two high-dimensional omics datasets, we utilize score tests and matrix calculations for computational efficiency. We further apply an empirical permutation method with genomic control to effectively control the family-wise error rate in studies with small sample sizes.

With real datasets, we demonstrate that *MITScan* achieves computational gains of three orders of magnitude compared to commonly used Wald or t tests, while identifying similar significant OTU-gene pairs. The *MITScan* R package is accessible on GitHub at https://github.com/yajing-hao/MITScan.

## 1 Introduction

The microbiome is dynamic and changes in response to environmental and physiological factors, including diet, stress, exercise, and antibiotics [Zheng et al., 2020]. It plays a critical role in maintaining homeostasis [Peterson et al., 2015], modulating immune responses [Shi et al., 2017, Jiao et al., 2020], and supporting metabolic functions [Cox et al., 2022]. Imbalances in the microbiome have been implicated in a wide range of various diseases, including inflammation [Belkaid and Hand, 2014], obesity [Maruvada et al., 2017], diabetes [Jamshidi et al., 2019], cardiovascular disease [Ahmadmehrabi and Tang, 2017], and oncogenesis [Scott et al., 2019]. Insights from microbiome research can inform therapeutic strategies to prevent and treat diseases by modulating the microbiome [Shukla et al., 2024].

Changes in the host transcriptome can influence microbial composition, while alterations in the microbiome can, in turn, modulate host gene expression, creating a dynamic and bidirectional relationship between the microbiome and host transcriptome [Nichols and Davenport, 2021]. Understanding these microbiome-host associations is essential for elucidating their roles in health and disease. Notably, genome-wide association studies (GWAS) have identified genetic variants that shape gut microbial diversity and demonstrated that host DNA plays a pivotal role in determining the abundance of specific bacterial taxa [Blekhman et al., 2015, Bonder et al., 2016, Davenport, 2016, Nichols and Davenport, 2021]. Similarly, quantitative trait loci (QTL) analyses have detected genomic regions associated with microbiome composition [Benson et al., 2010]. These findings suggest that host genetic regulation of the microbiome may be mediated through transcriptional mechanisms, as the host genome sequence remains unchanged despite the presence of microbiome [Nichols and Davenport, 2021]. Building on this foundation, our research aims to investigate how host-derived genes regulate microbial populations genome-widely, ultimately leading to differential microbial growth and community composition.

With the development of high-throughput sequencing technologies, 16S ribosomal RNA (rRNA) sequencing and internal transcribed spacer (ITS) sequencing have become common methods for identifying bacterial and fungal strains in the microbiome, respectively. Microbiome profiles are typically summarized as count matrices, where each entry representing the number of reads mapped to a given operational taxonomic unit (OTU) in a sample [He et al., 2015]. Meanwhile, host-targeted RNA-Seq is used to quantify gene expression levels in the host transcriptome. Joint analysis of the microbiome and host transcriptome data enables systematic investigations of their associations, revealing their impact on health and their contribution to the development of various disorders. For example, such associations have been explored in inflammatory bowel disease (IBD) [Morgan et al., 2015, Häsler et al., 2017, Lloyd-Price et al., 2019, Priya et al., 2022], colorectal cancer [Priya et al., 2022], non-small cell lung cancer [Zhou et al., 2023], and HBV-related hepatocellular carcinoma [Huang et al., 2020]. These studies have provided valuable insights into how genetic and microbial factors contribute to disease development and progression.

To identify associations between the microbiome and host transcriptome, most existing analyses often rely on Pearson correlation [Huang et al., 2020] or more robust Spearman correlation [Lloyd-Price et al., 2019]. However, microbiome count data can be extremely sparse with up to 90% zeros, resulting in zero inflation and overdispersion [Lin and Peddada, 2020]. These zeros may arise not only from biological absence but also from technical artifacts, such as insufficient sequencing depth, where taxa with low abundances are randomly missed entirely [Lutz et al., 2022]. Conventional Pearson and Spearman correlations between multi-omics datasets are susceptible to false positives due to the sparsity of microbiome data [Knight et al., 2018], limiting their utility for identifying true associations. More advanced statistical methods, including multivariate linear models [Morgan et al., 2015] and LASSO (least absolute shrinkage and selection operator) [Priya et al., 2022], have been proposed. Nonetheless, all these methods generally fail to adequately account for the fact that microbiome data are sparse count data with extra zeros, potentially yielding biased or inefficient estimates. Thus, there is a pressing need for statistical frameworks specifically designed to handle the sparsity and discrete nature inherent to microbiome datasets.

What’s more, 16S rRNA sequencing typically yields hundreds to thousands of OTUs, depending on how sequences are clustered at the genus or species level. And host-targeted RNA-Seq data generally captures expression profiles for over 20,000 genes. This leads to approximately 20 million (1,000 OTUs × 20,000 genes) potential association tests, highlighting the need for a computationally efficient analysis method, which is currently lacking.

To overcome these challenges, we present *MITScan*, a computationally efficient genome-wide **MI**crobiome-host **T**ranscriptome **Sc**ore-based association **an**alysis, extending *PETScan* with negative binomial models [Hao et al., 2025]. *MITScan* employs zero-inflated negative binomial (ZINB) models [Zhang et al., 2016, Risso et al., 2018] to accommodate extra zeros commonly observed in microbiome data, providing a more accurate and robust framework for studying how the microbiome is regulated by host gene expression. To address computational issues, *MITScan* implements two strategies to ensure its practical application. First, *MITScan* utilizes score tests based solely on parameter estimates under the null hypothesis. This approach is particularly efficient because, for a given operational taxonomic unit (OTU), its restricted estimates under the null hypothesis remain the same across all tested host genes and need to be computed just once. Second, *MITScan* boosts computational efficiency by leveraging efficient large matrix operations. In addition, the asymptotic *χ*^2^ distributions of score tests may not be valid when dealing with data from studies of limited sample sizes. To maintain control over family-wise error rates or false discovery rates in such settings, *MITScan* integrates an empirical permutation procedure inspired by genomic control [Devlin and Roeder, 1999, Zou et al., 2014].

We demonstrate the effectiveness of *MITScan* using the inflammatory bowel disease (IBD) multi-omics data from the Integrative Human Microbiome Project (iHMP) [Lloyd-Price et al., 2019]. This framework allows us to detect genome-wide regulatory effects of host genes on microbial abundances. Our findings reveal that associations between the microbiome and host genes contribute to dysregulation in IBD, highlighting the importance of integrative multi-omics approaches in deciphering complex disease mechanisms.

## 2 Methods

Consider a dataset of *N* paired samples of microbiome and host transcriptome data with *T* OTUs and *G* host genes, respectively. For OTU *t* (*t* = 1, …, *T*) and host gene *g* (*g* = 1, …, *G*), we define *y*_*ti*_ as the OTU occurrences and *w*_*gi*_ as the log-transformed gene expression of the *i*th sample (*i* = 1, …, *N*). For notational convenience, we omit subscripts *t* and *g* in subsequent discussions when focusing on the association between the OTU *t* and host gene *g* pair, provided the context is clear. To properly account for zero inflation in the microbiome count data *y*_*i*_, we apply a zero-inflated negative binomial model, where the count is zero with probability *π*_*i*_, whereas the count follows a negative binomial distribution with mean *µ*_*i*_ and dispersion parameter *ϕ* with probability 1 *− π*_*i*_. Let *x*_1*i*_, *x*_2*i*_, …, *x*_(*k−*1)*i*_ denote covariates related to the count component, while *z*_1*i*_, *z*_2*i*_, …, *z*_*mi*_ correspond to covariates for the zero component. Specifically,

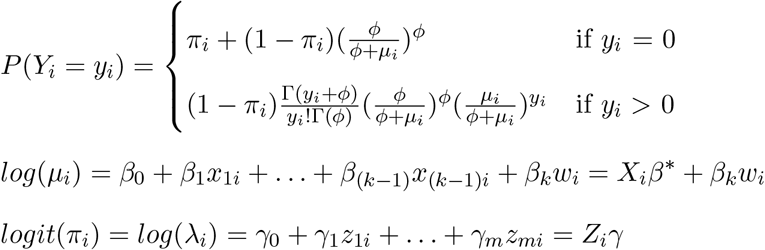

where 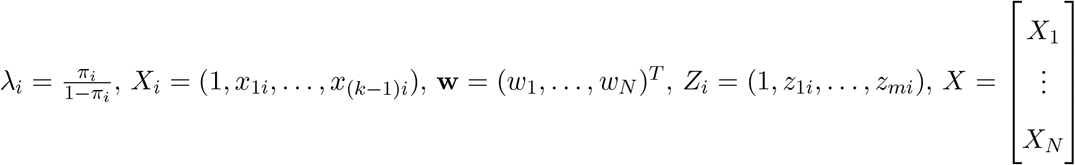

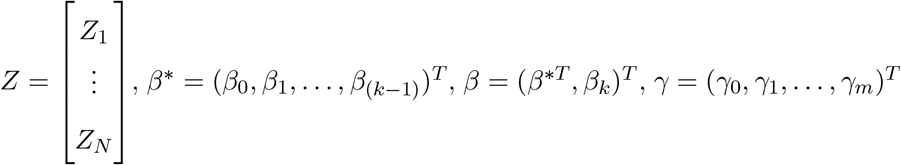 and θ = (ϕ, γ^*T*^, β^*T*^)^*T*^.

The log likelihood function of the data is therefore

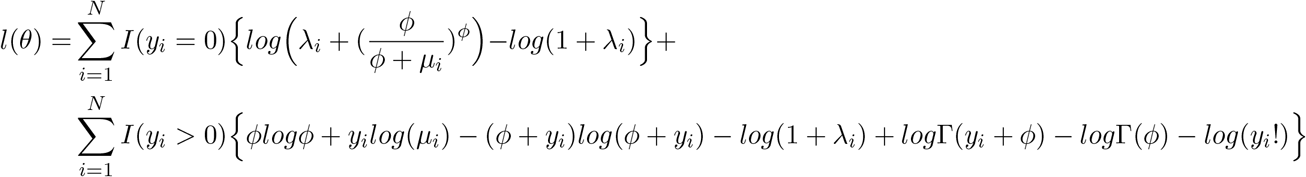

with the corresponding score function 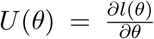 and the Fisher information matrix 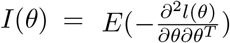. For the OTU and host gene pair, our primary interest is to test *H*_*0*_ : *β*_*k*_ = 0 versus *H*_*A*_ : *β*_*k*_ ≠ 0, determining whether the host gene expression influences the OTU abundances. Let 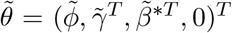 represent the restricted maximum likelihood estimate of *θ* under the null hypothesis. Importantly, this estimate 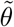 is independent of the host gene expression variable *w*_*gi*_. As a result, for OTU *t*, 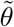 remains the same across all *G* host genes and only needs to be computed once.

To evaluate the null hypothesis *H*_0_ : *β*_*k*_ = 0, *MITScan* employs a score test, calculated as 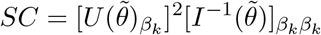 which follows 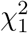 under *H*_0_ asymptotically, where 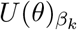 represents the (*m* + *k* + 3)th component of *U* (*θ*), and 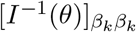 refers to the (*m* + *k* + 3, *m* + *k* + 3)th element of the inverse of *I*(*θ*), both corresponding to the parameter *β*_*k*_.Specifically, 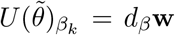, where 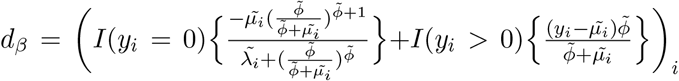 approximate the information matrix at 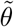 as

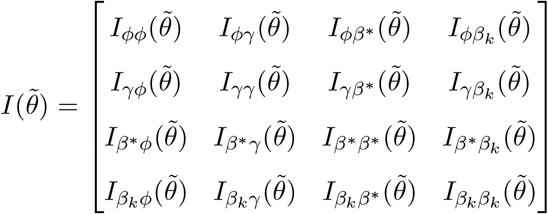

where 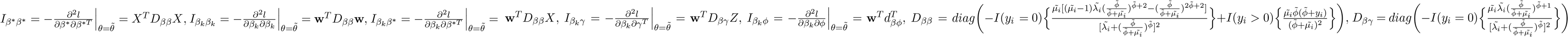 and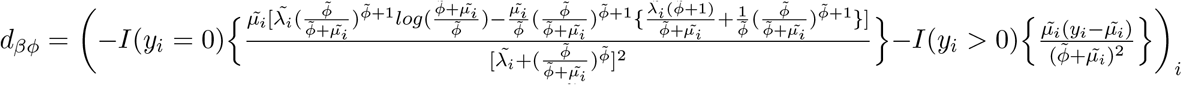.

Other terms are defined analogously and we omit 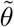 for notation simplicity. Notably, we observe that for a given OTU, 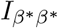 remains unchanged across all host genes and can be obtained from the null model, the same as 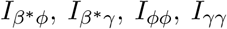, and *I*_*ϕγ*_. Consequently, we can avoid repeatedly calculating the upper-left (*m* + *k* + 2) *×* (*m* + *k* + 2) sub-matrix of the information matrix, that is,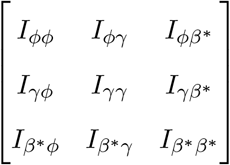, which we denote by *A* for notational convenience in the subsequent discussion, significantly enhancing computational efficiency. To further streamline computation, we leverage blockwise matrix inversion to extract specific elements of the inverse of Fisher information matrix without computing the full inverse of a large matrix, and eventually have

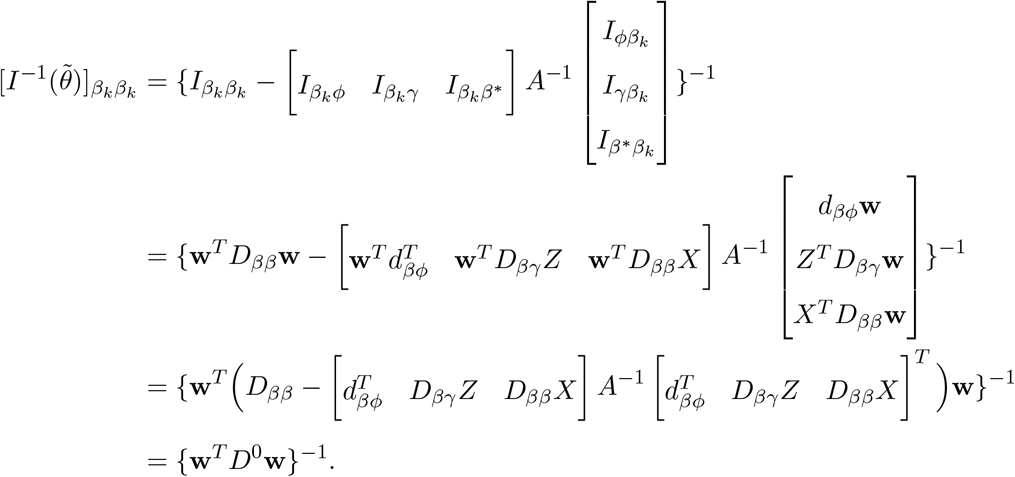

Again, for OTU *t, d*_*β*_, *D*_*ββ*_, *D*_*βγ*_, and *d*_*βϕ*_ all depend only on 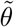 and therefore remain invariant across all *G* host genes. Moreover, since the sub-matrix *A* is independent of host genes, its inverse needs to be computed only once per OTU. As a result, the score test for one host gene can be extended easily to all host genes using matrix calculations as follows. Let **W** = [**w**_1_, …, **w**_*G*_], then **SC** = (*SC*_1_, …, *SC*_*G*_) = (*d*_*β*_**W**) ⊙ [*diag*(**W**^*T*^ *D*^0^**W**)]^ℜ^ ⊙ (*d*_*β*_**W**), where ⊙ represents the Hadamard (element-wise) product, *B*^ℜ^ denotes the element-wise reciprocal of a vector or matrix *B*, and 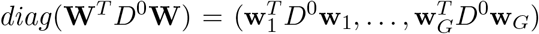. In contrast to the Wald and likelihood ratio tests, score tests for a given OTU necessitate modeling the zero-inflated negative binomial model under the null hypothesis just once for all host genes, rather than *G* full models separately for each host gene. This feature enables the proposed matrix operations to compute the score tests for all host genes simultaneously, avoiding repeated calculations for one host gene at a time, ensuring efficient computation.

For studies with limited sample sizes, score tests may exhibit inflation, compromising the validity of p-values derived from the asymptotic 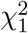 distribution. To ensure robust inference, we adopt an empirical permutation method with genomic control [Devlin and Roeder, 1999] to address this issue. For OTU *t*, let the observed score test statistics across the *G* host genes be represented by *SC* = (*SC*_1_, …, *SC*_*G*_). We then randomly sample the host transcriptome data without replacement *L* times, generating the corresponding permuted score tests as 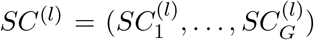, where *l* = 1, …, *L*. Using the genomic control framework, we empirically estimate the inflation factor *λ* as 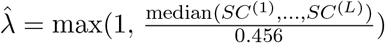 and rescale the observed score test statistics *SCg* to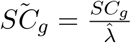 for *g* = 1, …, *G*, which are then compared to 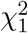 for p-value calculations.

## 3 Results

### 3.1 Simulation

To evaluate the performance of score-based association analysis using ZINB models, we generate synthetic microbiome and host transcriptome data. Under the null hypothesis of no association between microbiome and host transcriptome data, we simulate *y*_*i*_ with *log*(*µ*_*i*_) = *β*_0_ + *β*_1_*x*_*i*_ and *log*(*λ*_*i*_) = *γ*_0_ + *γ*_1_*z*_*i*_, where *i* = 1, …, *N*. Let *β*_0_ = 0, *β*_1_ = 1, *γ*_1_ = 0.2, *x* follow a normal distribution *N* (1, 1), and *z* follow *N* (0, 1). Then we generate *w* from *N* (2, 1), simulating the log-transformed expression level for a host gene. The Wald, score, and likelihood ratio tests are applied to determine whether *w* should be included in the model. We vary *γ*_0_ = −3, −2, −1, −0.5, 0, corresponding to different proportions of zeros (about 5%, 12%, 27%, 38%, 50%, respectively), thedispersion parameter *ϕ* = 5, 10, 20, and the sample size *N* = 50, 100, 200, 500, 1000. Each scenario is simulated over 10,000 repetitions to evaluate the empirical type I error rates. In addition, under the alternative hypothesis assuming there are associations between microbiome and host transcriptome data, we simulate *y*_*i*_ with *log*(*µ*_*i*_) = *β*_0_ + *β*_1_*x*_*i*_ + *β*_2_*w*_*i*_ and *log*(*λ*_*i*_) = *γ*_0_ + *γ*_1_*z*_*i*_. We vary *β*_2_ to 0.1, 0.2, and 0.3 to evaluate the empirical power of score tests. All other parameters are kept unchanged as described above.

Figure 1 presented the empirical type I error rates of the score, Wald, and likelihood ratio tests for ZINB models by sample sizes, proportions of zeros, and dispersion parameters. The results demonstrated that all three tests effectively controlled type I error when the sample size was 500 or larger. However, none of the three tests provided adequate type I error control in datasets with higher proportions of zeros and greater dispersion (smaller *ϕ*). Among the three methods, the score test was the least conservative, whereas the likelihood ratio test proved to be the most conservative. Additionally, differences among three tests diminished as sample sizes increased. All three tests controlled type I error better with smaller sample sizes after applying genomic control methods (Figure 2).

**Figure 1:**
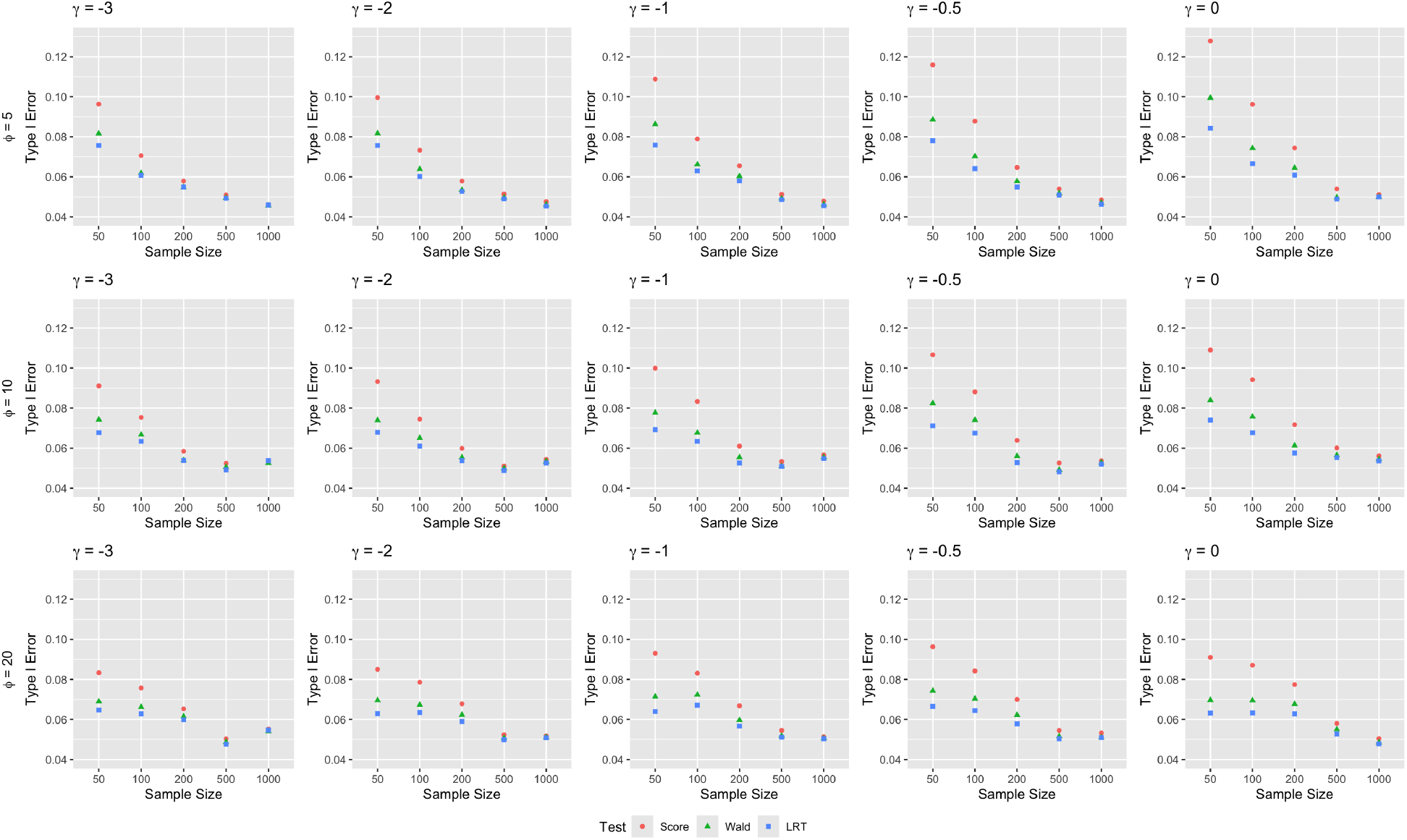
Empirical type I error rates of the score, Wald, and likelihood ratio tests for zero-inflated negative binomial models by sample sizes, proportions of zeros, and dispersion parameters.

**Figure 2:**
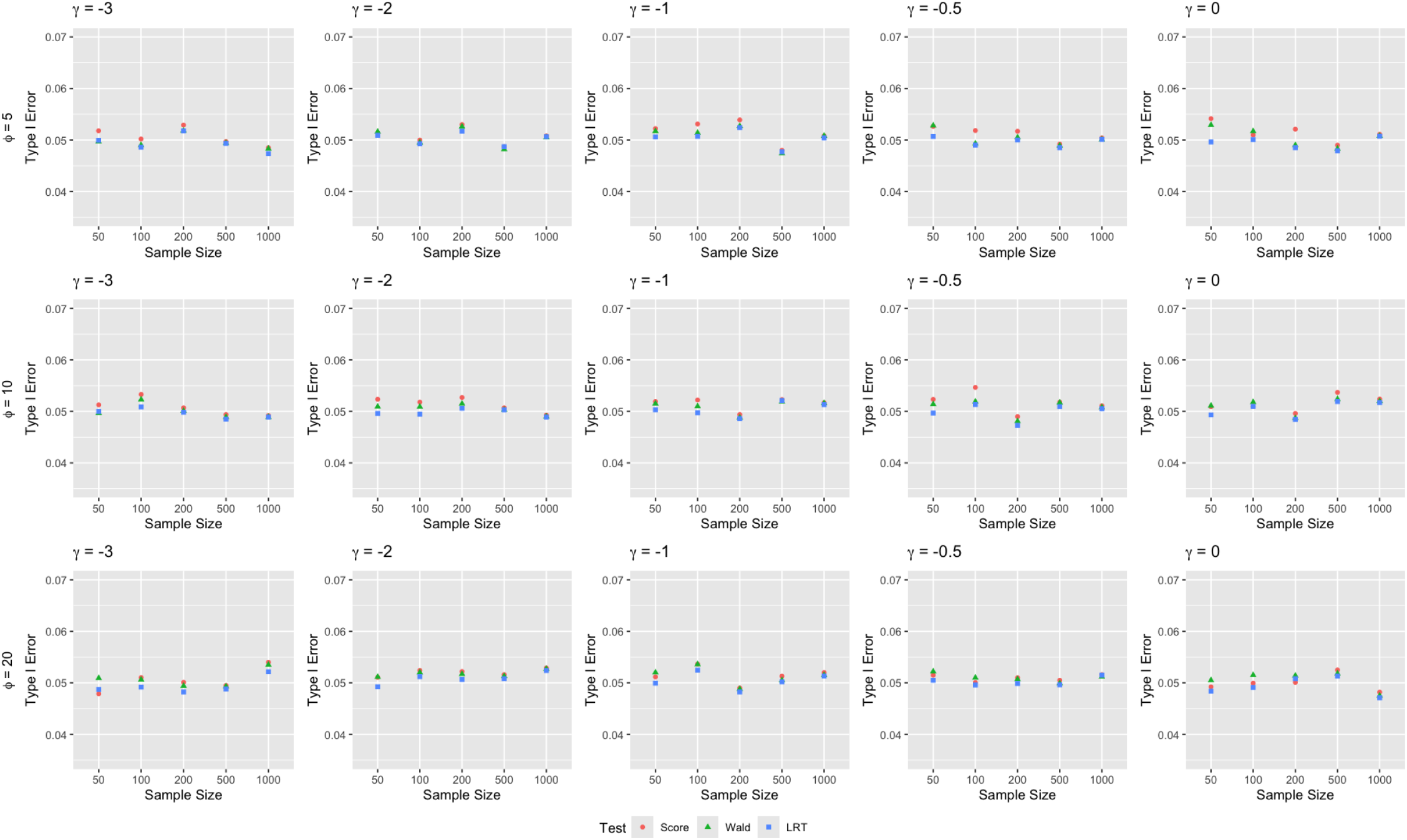
Empirical type I error rates of the score, Wald, and likelihood ratio tests for zero-inflated negative binomial models after applying genomic control by sample sizes, proportions of zeros, and dispersion parameters.

Figure 3 illustrated the empirical power of the score, Wald, and likelihood ratio tests at the 0.05 significance level for ZINB models by effect sizes, sample sizes, and proportions of zeros, with dispersion parameter *ϕ* = 10. The findings indicated that all three tests exhibited comparable power, which increased with larger effect sizes and sample sizes. However, in datasets with a high proportion of zero counts, statistical power remained insufficient when the sample size was small, even with a large effect size.

**Figure 3:**
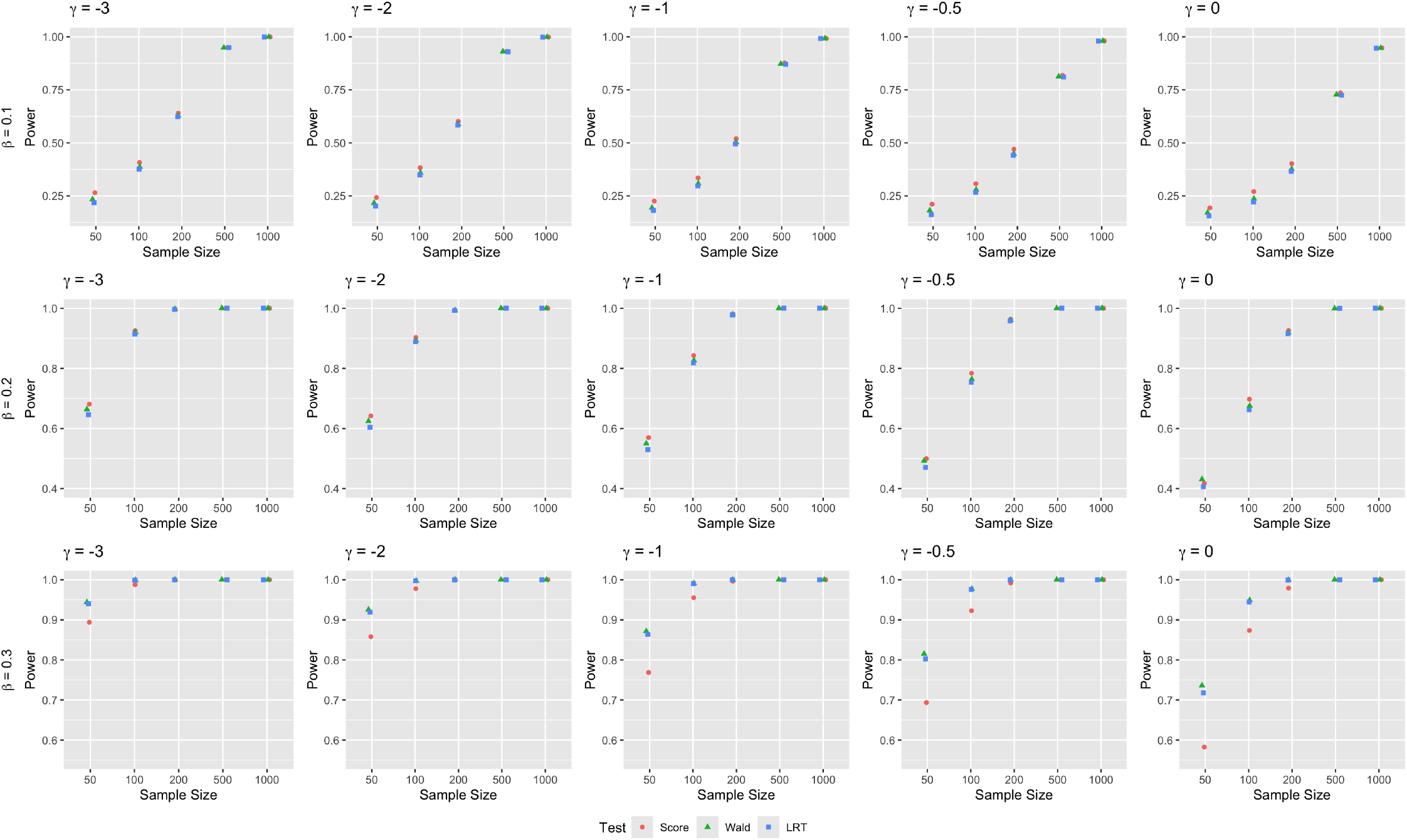
Empirical power of the score, Wald, and likelihood ratio tests for zero-inflated negative binomial models after applying genomic control methods by effect sizes, sample sizes, and proportions of zeros with dispersion parameter *ϕ* = 10.

### 3.2 Microbiome and Host Transcriptome

Inflammatory bowel diseases (IBD), including Crohn’s disease (CD) and ulcerative colitis (UC), are marked by chronic relapsing and remitting inflammation of the gastrointestinal tract or colon, often resulting in significant debilitation. The Integrative Human Microbiome Project (iHMP) collected a comprehensive multi-omics dataset on IBD, known as the inflammatory bowel disease multi-omics data (IBDMDB) [Lloyd-Price et al., 2019]. They followed 132 participants for one year and collected stool, biopsy, and blood specimens for longitudinal host and microbial activity.

Biopsies were profiled using 16S rRNA sequencing for microbial taxa and RNA-Seq for host gene expression. If multiple time points were available for an individual in paired datasets, only the data from the earliest week were retained. Data were downloaded from https://ibdmdb.org/results. In their study, associations between microbial taxonomic profiles and host gene expression were evaluated using partial Spearman correlation, adjusting for BMI, age at consent, sex, and IBD diagnosis, with analyses conducted separately for rectal and ileal tissues [Lloyd-Price et al., 2019]. We repeated their analysis using *MITScan*. For the rectum, 64 samples had paired microbial taxonomic profiles and host gene expression data. After filtering out OTUs with counts greater than 5 in ≤ 3 samples and genes with counts greater than 10 in ≤ 3 samples, 181 OTUs and 21,849 genes remained, resulting in 3,954,669 OTU-gene pairs tested for association, among which 7,757 significant OTU-gene pairs were detected by *MITScan* with Benjamini & Hochberg (BH) adjusted p-values less than 0.05. The analysis completed in less than 75 seconds when parallelized across 64 cores, demonstrating a significant computational improvement over conventional Wald tests, which were estimated to require approximately 1,000 times longer.

Some of the genes associated with the significant OTU-gene pairs are known to play a role in the development of IBD (Figure 4). For instance, *NOD2* is the first susceptibility gene to Crohn’s disease discovered at the end of the 20th century [Hugot et al., 2001], and its genetic variants are associated with the stricturing phenotype, ileal involvement, and an increased risk of steroid refractoriness [Kayali et al., 2024]. We found that *NOD2* was associated with the abundance of *Ruminococcus 1* and *Ruminococcaceae UCG 013*. Furthermore, recent evidence has emphasized immune system dysfunction, particularly toll-like receptor (TLR)-mediated innate immune dysfunction, as key factors in the pathogenesis of IBD [Lu et al., 2018]. Specifically, *TLR3* and *TLR7* promote protective immunity under inflammatory conditions, while *TLR4* contributes to intestinal tissue destruction and ulceration, and *TLR8* induces mucosal inflammation [Junker et al., 2012, Sainathan et al., 2012, Saruta et al., 2009]. Within the *TLR* family, we identified correlations between *TLR3, TLR4, TLR7, TLR8*, and *TLR10* with the abundance of *Fusicatenibacter, Roseburia, Odoribacter, Citrobacter*, and *Aggregatibacter*, respectively. Besides, *IL10RA* and *IL10RB*, which encode the IL10R1 and IL10R2 proteins, respectively, form a heterotetramer that constitutes the interleukin-10 receptor. Glocker’s findings demonstrated that loss-of-function mutations in either *IL10RA* or *IL10RB* are present in children with severe early-onset enterocolitis and the most prominent phenotype in patients with IL10R1 or IL10R2 deficiencies is IBD [Glocker et al., 2009].

**Figure 4:**
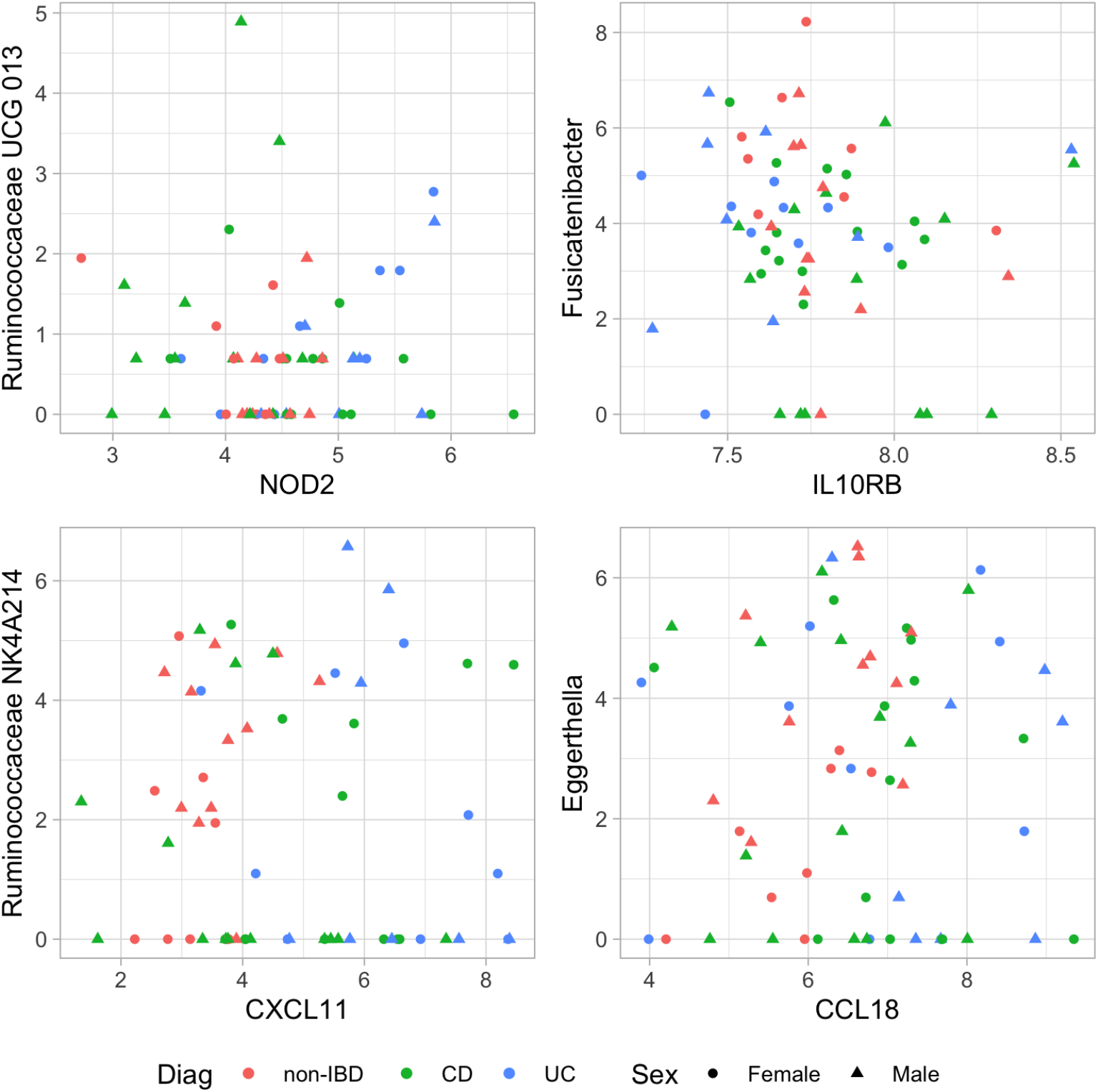
Scatter plots of significant OTU-gene pairs identified in the rectum.

We showed that *IL10RB* was correlated with the abundance of *Fusicatenibacter* in the rectum. Our results suggested that specific microbial taxa may be influenced by host genes, potentially contributing to IBD progression. This strengthened the biological significance of our analysis and provided insights into potential therapeutic strategies aimed at modulating the gut microbiome through host-targeted interventions.

In addition, chemokines play a critical role in recruiting leukocytes to sites of inflammation and infection and have also demonstrated direct antimicrobial activity in vitro [Murphy, 2019]. They are categorized based on the number and arrangement of conserved cysteine residues [Murphy et al., 2000]. In humans, CC chemokines feature two adjacent conserved cysteines, whereas in CXC chemokines the first two conserved cysteines are separated by one amino acid [Yung and Murphy, 2012]. Specifically, *CCL3* and *CCL3L1* recruits and activates macrophages and neutrophils [Bonville et al., 2009], while *CCL4* and *CCL4L1* primarily recruits macrophages and monocytes [Lu et al., 2004]. *CCL18* is primarily expressed by dendritic cells and macrophages [Schutyser et al., 2005]. Although their direct antimicrobial properties are not well-documented, these chemokines indirectly contribute to antimicrobial defense by promoting inflammation and enhancing immune cell activity. What’s more, *CXCL10* has been shown to directly kill a wide variety of pathogenic bacteria in addition to its receptor-dependent role in regulating immune-cell migration and activation [Crawford et al., 2023]. *CXCL14* has also demonstrated a broad range of antimicrobial effects [Lu et al., 2016]. Consistent with this, previous studies have shown that antimicrobial genes exhibited higher expression in patients with IBD compared to healthy controls [Lloyd-Price et al., 2019, Bennet et al., 2018]. This increased expression may contribute to the microbiome dysbiosis observed in IBD, as host-derived antimicrobial factors can directly alter gut microbial composition [Nichols and Davenport, 2021]. Significant OTU-gene pairs associated with chemokines identified in the rectum were shown in Table 1. The indirect or direct antimicrobial effects of chemokines aligned with our findings, as they suggested that the chemokine-related genes may contribute to maintaining homeostasis by regulating microbial communities.

**Table 1:** Significant OTU-gene pairs associated with chemokines identified in the rectum.

| Gene | OTU | FDR | Gene | OTU | FDR |
| --- | --- | --- | --- | --- | --- |
| CCL3 | Subdoligranulum | 0.0355 | CXCL10 | Roseburia | 0.0431 |
| CCL3L1 | Ruminococcus 1 | 0.0082 | CXCL11 | Roseburia | 0.0002 |
| CCL4 | Subdoligranulum | 0.0391 | CXCL11 | Ruminococcaceae NK4A214 | 0.0413 |
| CCL4L1 | Aggregatibacter | 0.0041 | CXCL12 | Fusicatenibacter | 0.0022 |
| CCL18 | Eggerthella | 0.0492 | CXCL14 | Roseburia | 0.0262 |
| CXCL5 | Ruminococcus 2 | 0.0299 | CXCL17 | Ruminiclostridium 5 | 0.0360 |

To further understand the genes associated with the significant OTU-gene pairs, we performed Kyoto Encyclopedia of Genes and Genomes (KEGG) enrichment analysis on these genes using the clusterProfiler R package [Yu et al., 2012]. The analysis revealed that they were enriched for neurodegenerative disease pathways, including Alzheimer’s disease (AD), Parkinson’s disease (PD), and prion diseases. Epidemiological studies have reported that patients with IBD exhibit a significantly increased risk of developing AD [Zhang et al., 2021,Wang et al., 2022,Zong et al., 2024] and PD [Brudek, 2019,Villumsen et al., 2019,Zong et al., 2024] compared to the general population, although the underlying mechanisms are not completely clear. One partial explanation is that the microbiota–gut–brain axis is a bidirectional communication network connecting the gastrointestinal tract with the central nervous system [Carabotti et al., 2015, Cryan et al., 2019, Wang et al., 2022, Kim et al., 2023, Loh et al., 2024]. These findings underscored the potential association between intestinal inflammation and neurodegenerative processes, highlighting the importance of further research into shared biological pathways and therapeutic targets.

Next, we analyzed the paired data from the ileum, which also comprised 64 participants. Applying the same filtering criteria as used for the rectal data, we retained 173 OTUs and 22,210 genes. Out of 3,842,330 tested OTU–gene pairs, *MITScan* identified 5,083 significant pairs with BH-adjusted p-values less than 0.05. Notably, several genes involved in these significant OTU–gene pairs have been previously reported as IBD-associated. Growing evidence has demonstrated a strong association between altered gut microbiota and decreased aryl hydrocarbon receptor (*AHR*) activity in IBD [Hou et al., 2023]. In healthy intestinal epithelial cells, *AHR* is typically highly expressed and activated [De Vos et al., 2022], whereas lower *AHR* levels have been detected in IBD patients [Monteleone et al., 2011]. Our analysis further revealed a correlation between *AHR* and the abundance of *Ruminococcaceae UCG 003*. In addition, Darnaud demonstrated that mice with hepatocytes expressing human *REG3A* exhibited increased resistance to colitis, and the fecal microbiota of these mice showed a significant compositional shift compared to control mice, with an enrichment of *Clostridiales* (*Ruminococcaceae, Lachnospiraceae*) and a reduction in *Bacteroidetes* (*Prevotellaceae*) [Darnaud et al., 2018]. In line with these findings, we found that *REG3A* was associated with the abundance of *Anaerostipes*, a member of the *Lachnospiraceae* family. This association suggested that *REG3A* may contribute to shaping the gut microbiome by promoting the growth of beneficial taxa, such as *Anaerostipes*. Moreover, *MUC2* encodes a protein in the mucin family, and its deficiency alters mucus composition, potentially influencing the onset or progression of IBD. In ulcerative colitis (UC), the mucus is observed to be thinner and discontinuous, with lower levels of MUC2 mucin detected during active phases [Liu et al., 2020]. We identified a correlation between *MUC2* and the abundance of *Intestinibacter*, suggesting that changes in mucin production may influence specific microbial populations in the gut, potentially contributing to disease pathogenesis. Several significant OTU-gene pairs identified in the ileum were summarized in Table 2, providing further evidence of the complex influence of host genes on the gut microbiome.

**Table 2:** Significant OTU-gene pairs identified in the ileum.

| Gene | OTU | FDR | Gene | OTU | FDR |
| --- | --- | --- | --- | --- | --- |
| AHR | Ruminococcaceae UCG 003 | 0.0390 | CXCL3 | Phascolarctobacterium | 0.0069 |
| CCL1 | Rothia | 0.0265 | CXCL11 | Lachnoclostridium | 0.0120 |
| CCL4L1 | Erysipelatoclostridium | 0.0111 | CXCL14 | Lachnospira | 0.0491 |
| CCL8 | Parvimonas | 0.0004 | MUC2 | Intestinibacter | 0.0157 |
| CCL14 | Ruminococcaceae UCG 002 | 0.0009 | REG3A | Anaerostipes | <0.0001 |
| CCL23 | Intestinibacter | 0.0084 | TLR3 | Haemophilus | 0.0003 |
| CXCL1 | Subdoligranulum | 0.0434 | TLR8 | Ruminococcaceae UCG 003 | 0.0298 |

## 4 Discussion

*MITScan* sheds light on the relationship between the microbiome and host transcriptome, providing valuable insights into how host gene regulates microbiome composition. With the IBD data, our analysis identified several significant OTU-gene pairs, with the associated genes playing key roles in IBD development, underscoring the biological relevance of our approach. By deepening our understanding of these complex associations, *MITScan* holds potential for guiding novel therapeutic strategies that target microbiome composition through modulation of host gene expression, paving the way for innovative approaches to disease prevention and treatment.

A major challenge in microbiome data analysis is the high prevalence of excess zeros, arising from both biological factors (the presence of low-abundance taxa) and technical limitations (sequencing depth constraints). Employing ZINB models, *MITScan* effectively handles sparsity and overdispersion commonly observed in microbiome data. However, complex ZINB models may fail to converge, particularly in datasets with limited sample sizes. In such cases, simpler models like zero-inflated Poisson (ZIP) models may be more appropriate. While ZIP models do not include a dispersion parameter and are inherently simpler than ZINB models, they retain the capacity to capture zero inflation, making them a viable alternative for sparse count data. *MITScan* implements ZIP models as an option.

Furthermore, *MITScan* is versatile and extends beyond paired microbiome and host transcriptome data. It is applicable to a broad range of paired high-dimensional datasets, as long as ZINB models provide an appropriate framework for data modeling. For instance, in metabolomics research, metabolite concentration data often exhibit similar zero-inflation patterns, for which *MITScan* is suitable for analyzing paired metabolomics-transcriptomics data [Cavill et al., 2016]. Likewise, in investigations of host-pathogen associations by dual RNA-Seq, transcriptomic profiles of host cells often contain sparse signals of pathogen-derived transcripts, making ZINB models an ideal choice for association analyses in infectious disease research [Westermann et al., 2017,Sounart et al., 2023]. This adaptability ensures that *MITScan* remains relevant across different fields of study, contributing to a deeper understanding of complex biological systems.

In most cases, the covariates used in the zero component are a subset of those included in the count component. However, our formula remains valid regardless of whether *Z* is fully contained within *X* or not. This flexibility ensures the robustness and broad applicability of our approach across different model specifications. Furthermore, while our current implementation of *MITScan* includes host gene expression levels only in the count component of the model, it can be easily extended to incorporate them into the zero component as well.

In addition, *MITScan* assesses the presence of associations without determining whether a host gene positively or negatively influences OTU abundances, due to its reliance on score tests. Despite this limitation, *MITScan* serves as a valuable screening tool for microbiome-host associations, providing a foundation for subsequent in-depth investigations. While our study primarily investigates how host genes influence the microbiome, the impact of the microbiome on host gene expression is also of interest, given the bidirectional nature of these interactions [Nichols and Davenport, 2021]. For such analyses, we recommend using *PETScan*, a computationally efficient genome-wide ***PE***ak-***T***ranscript ***Sc***ore-based association ***an***alysis, developed by our group to test associations of RNA-Seq and other high-dimensional omics data. *PETScan* employs negative binomial models specifically designed for RNA-Seq data, enabling robust and accurate analyses [Hao et al., 2025].

In summary, *MITScan* offers a computationally efficient and biologically meaningful framework for investigating genome-wide microbiome-host associations, providing a deeper understanding of host-microbiome crosstalk in health and disease.

